# PRISM: A platform for illuminating viral dark matter

**DOI:** 10.1101/2025.07.24.666652

**Authors:** Han Zhang, Justin Boeckman, Rohit Gupte, Ashlee Prejean, Jonathan Miller, Alexandra Rodier, Paul de Figueiredo, Mei Liu, Jason Gill, Arum Han

## Abstract

The exploration of bacteriophage diversity remains constrained by reliance on conventional plaque assays, which bias discovery toward phages that form visible plaques and propagate efficiently. The discovery process, as well as measuring correct phage concentrations and phage resistance frequencies, are also time-consuming and labor intensive. Here, we present PRISM (Phage Recovery and Investigation in Single-droplet Microenvironments), a droplet microfluidics-based, plating-independent platform for the high-throughput isolation, quantification, and characterization of bacteriophages. By encapsulating phage–host interactions in water-in-oil microdroplets at single-phase resolution, and then coupling them with fluorescence-based detection and sorting at million-droplet scale, PRISM enables rapid identification of novel phages, including those that evade detection in conventional assays. We demonstrate PRISM’s capabilities across several key applications: (i) recovery of both novel “plaquing” and “non-plaquing” *Salmonella* phages from a concentrated environmental sample, where PRISM recovered multiple representatives of a non-plaquing genus completely missed by conventional plating-based approaches; (ii) accurate titering of poor-plaquing phages such as *Rhodococcus* phage ReqiDocB7, for which PRISM reported a 2-fold higher titer than conventional plaque assays; and (iii) plating-free determination of phage resistance frequency, with PRISM yielding resistance and lysogeny frequencies (e.g., 1 in 4.4 × 10^5^ cells for T7 *Escherichia* phage resistance and 1 in 21 cells for *Escherichia* Lambda phage lysogeny) that closely match conventional estimates. These data highlight PRISM’s enhanced resolution and accuracy, as well as its ability to detect rare, non-plaquing, or slow-replicating phages, in a high speed and automated fashion, thereby underscoring the platform’s power to access viral “dark matter” traditionally overlooked using conventional methods.

## Introduction

Bacteriophages, or phages, are viruses that infect bacteria and have gained renewed interest for their therapeutic potential and industrial applications in the post-antibiotic era^1^. Phages are also known to be important players in microbial ecosystems, driving bacterial cell turnover and the global carbon cycle by inducing host cell lysis and redirecting nutrients^2 3^. The advent of modern sequencing technologies, such as whole genome sequencing (WGS) and metagenomic sequencing, have profoundly impacted phage research and led to a substantial surge in the identification of new phage genomes. These advances also revealed extensive phage diversity and a large reservoir of unknown and uncharacterized genes^4^. However, most phages identified through metagenomics remain uncultured due to limitations imposed by traditional phage culture methods^5^ and the absence of an approach that enables rapid and high throughput isolation and amplification.

Methods for isolating and characterizing phages from the environment have remained largely unchanged for more than 100 years^6,7^. Conventional phage isolation methods have several limitations (**Fig. 1A, top**). The process is time- and labor-intensive, often depending on liquid enrichment and multiple rounds of single-plaque isolation on soft agar overlays for phage purification^8,9^. This isolation from enrichments of mixed phage populations is often biased towards phages with an initial higher abundance and/or better propagation characteristics under standard laboratory conditions, which rely on the lytic activity of phages to form zones of clearing, or plaques, on bacterial lawns. This approach, therefore, results in a constrained representation of phage diversity, overlooking phages that are slow to replicate. It also fails to capture phages that are unable to form visible plaques in soft agar. These challenges necessitate plating-independent universal phage isolation approaches that enable the capture of single interaction events between host cells and phages and that facilitate the isolation of undiscovered phages through discrete parallel propagation of mixed phage populations.

**Fig. 1.**
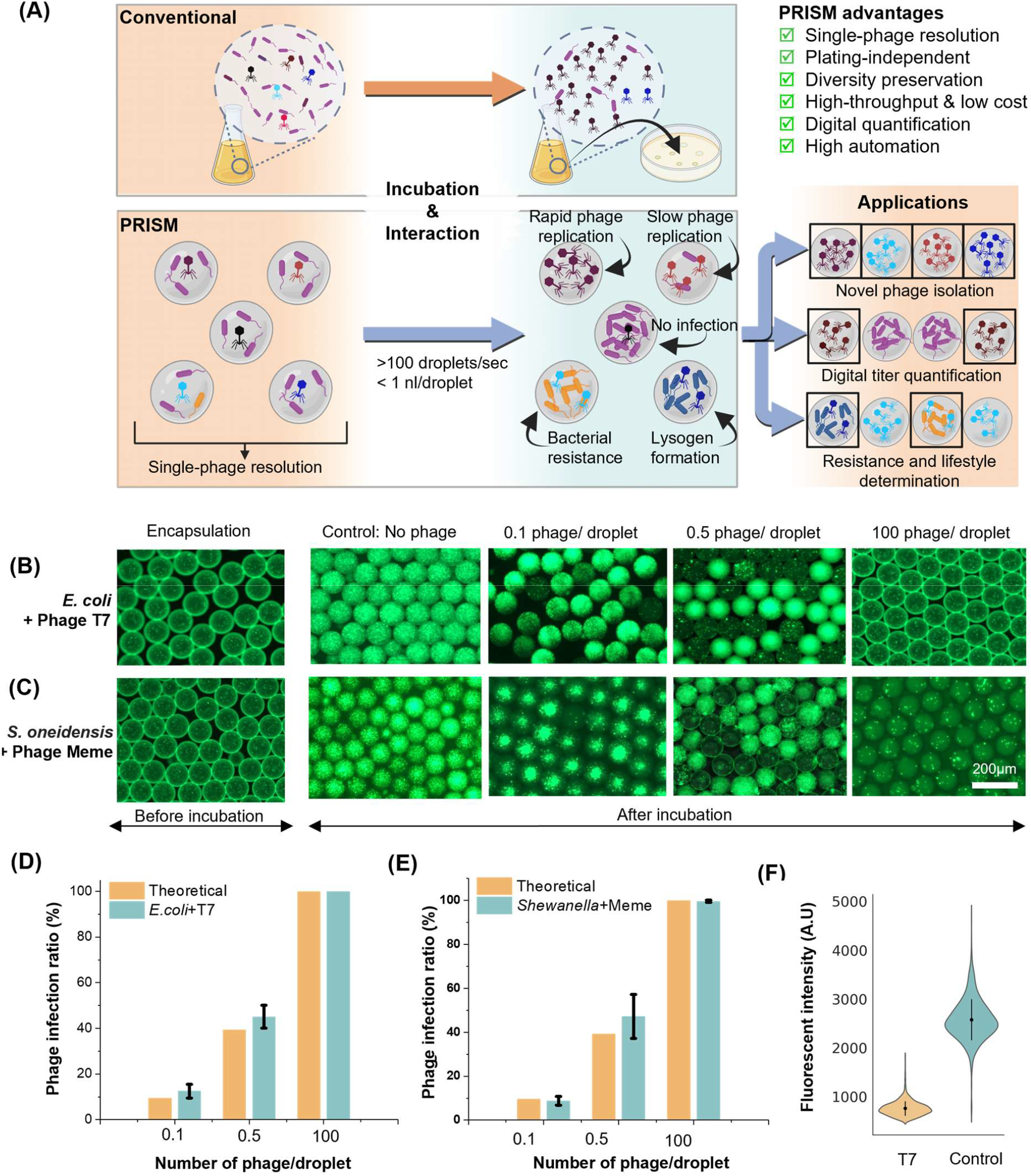
Workflow of PRISM. **(A)** Comparison of conventional plaque assay and the droplet microfluidics-based PRISM workflow. **(B–C)** Bacterial killing in droplets by species-specific phages: T7 for *E. coli* and Meme for *S. oneidensis*, tested at three phage concentrations (phages per droplet). Here 10-20 bacteria cells were initially encapsulated in the droplets. **(D–E)** Observed infection ratios for *E. coli* + T7 **(D)** and *S. oneidensis* + Meme **(E)** aligns with the expected Poisson distribution of phage encapsulation, supporting single-phage resolution (N = 3). **(F)** Fluorescence intensity of *E. coli* droplets with and without T7 phage; lower intensity indicates bacterial inhibition, enabling sorting of active (low-intensity) vs inactive (high-intensity) phage-containing droplets.

Following isolation, phages are evaluated for therapeutic or industrial potential based on their ability to lyse host bacteria. An additional consideration for applied use is phage lifestyle: virulent phages are restricted to rapidly killing hosts through the lytic pathway, while temperate phages can also integrate into the host genome as dormant prophages (lysogeny)^1,7^. Temperate phages are generally avoided in therapy due to rapid formation of lysogenized, phage-insensitive bacteria. The frequency of lysogeny depends on phage-specific factors and can increase with high multiplicity of infection (MOI)^10,11^, or, if MOI independent, remain constant at rates much higher than spontaneous host resistance. Phage lifestyle as well as resistance frequency are key screening criteria for determining the applied potential of a given phage. However, they are traditionally assessed via plating-dependent agar-based assays that are low throughput and difficult to automate^11-13^.

Droplet microfluidics has been shown to be a powerful tool for microbiology applications, enabling single-cell encapsulation and the rapid screening of hundreds of thousands of individual cell-encapsulated droplets^14-16^. As a complementary and advantageous alternative to traditional methods, we describe here the development and validation of versatile microdroplet-based phage techniques (**Fig. 1A**), which provide a streamlined, universal, and plating-independent approach for phage biology, including isolation, quantification, lifestyle prediction, and resistant mutant identification. By encapsulating host bacteria and phages into water-in-oil microdroplets at single-phase resolution, numerous spatially distinct “micro-enrichments” are created, facilitating rapid and clonal phage isolation while preserving diversity and eliminating the need for the optimization of conditions for time-consuming plaque formation. We term this technology “**PRISM”** (**P**hage **R**ecovery and **I**nvestigation in **S**ingle-droplet **M**icroenvironments). By altering phage to host ratios as well as “hit” droplet criteria, in addition to isolation, we demonstrate that PRISM can also be used for a variety of plate-independent phage characterization applications.

## Results

### Identification of bacterial growth inhibition with single-phage resolution

To demonstrate that phage-mediated host killing can be quantified using PRISM, we utilized the well-characterized model *Escherichia* phage T7 (**Fig. 1B**) and the recently isolated *Shewanella oneidensis* phage Meme (**Fig. 1C**). Various concentrations of phage were combined with exponentially growing host cells and a fluorescent cell viability dye SynaptoGreen™ C4 (Biotium) (for quantification of live cells in droplets) immediately before droplet generation. After incubation, microdroplets were screened for bacterial growth based on fluorescence intensity and confirmed by direct visualization under brightfield and fluorescent microscopy. High fluorescence intensity indicates bacterial growth and the absence of host cell-affecting phage in a droplet (**Fig. 1B and 1C**). Conversely, low fluorescence intensity indicated low bacterial growth, signifying the presence of phage that inhibited growth of the target host within the droplet. With approximately 0.1, 0.5, and 100 T7 phage particles per droplet (diluted using plaque forming unit [PFU]/mL titer determined by conventional plaque-assay), the percentage of low-intensity droplets was, as expected, 12.5 ± 2.9 (standard deviation, SD), 45.8 ± 5.1 (SD), and 100 ± 0 (SD), respectively. Similarly, when phage Meme was similarly tested at three phage concentrations (**Fig. 1D**), the corresponding percentages of low-intensity droplets were 8.8 ± 2.0 (SD), 47.2 ± 9.8 (SD), and 99.6 ± 0.004 (SD), respectively. In both cases, approximately 10 – 20 bacterial cells were initially encapsulated per droplet (**Fig. S1)**. These percentages were close to the expected values based on Poisson distribution^17^. Increasing the phage concentration to approximately 100 phages per droplet resulted in near 100% growth inhibition, as indicated by uniformly low fluorescence intensity across droplets (**Fig. 1D & E**). These results confirm that PRISM enables quantitative, single-phage-resolution detection of bacterial growth inhibition in droplets.

Next, to demonstrate the high-throughput screening potential of PRISM, we leveraged the inverse correlation between phage activity and fluorescence to distinguish droplets with and without phage using optical laser signal detection (**Fig. 1F**). Droplets containing T7 phage showed markedly lower fluorescence intensity (mean intensity = 750 ±150, *N*=*1426*) due to inhibited *E. coli* growth, while control droplets (no phage) retained high fluorescence intensity (mean intensity = 2565 ± 429, *N=910*) from higher numbers of bacteria due to growth. This binary fluorescence readout enabled digital, single-phage-resolution screening of functional phage–host interactions at rates greater than 100 droplets/second.

### PRISM isolation of plaquing and non-plaquing phages

To demonstrate the utility of PRISM, we isolated novel phages infecting a clinically relevant strain of *Salmonella enterica* serovar Typhimurium (*S*. Typhimurium ATCC 53648) that constitutively expressed red fluorescent protein (RFP). Briefly, filter-sterilized wastewater samples were pooled and virions concentrated via PEG precipitation. This concentration process employing physical and chemical methodologies was selected to preserve phage diversity and avoid “winner take all” phage-host propagation dynamics^6,18,19^. Phage-encapsulated microdroplets generated from this sample were incubated overnight before laser-based detection and sorting for droplets in which phage-mediated growth inhibition of host cells (*S*. Typhimurium) could be detected (**Fig. 2A**). Notably, input phage concentration was kept sufficiently low such that each microdroplet contained no more than a single phage particle. Representative droplet images of key steps in the PRISM workflow are shown in **Fig. 2A**, including (1) rare droplets in which phage inhibited bacterial cell growth (RFP-dark) surrounded by droplets containing high-densities of bacteria (RFP-bright) in the initial library (**Fig. 2A Input**), (2) sorted droplets enriched for phage that inhibited bacteria growth (**Fig. 2A Sorting & Hits**), and (3) sorted droplets dispensed onto top-agar overlays for infectivity validation. Infectivity was confirmed by the presence of plaque formation and/or zones of clearance from the dispensed RFP-dark droplets on *Salmonella* top-agar lawns (**Fig. 2A Dispensing**). In parallel, to benchmark our findings we performed phage hunts using conventional approaches from the same PEG-concentrated sample (**Fig. 2A, top)**. Utilizing genetic sequences of these conventionally isolated phages, PCR primers were designed to de-duplicate PRISM-identified phages. Unique PRISM phage isolates were then sequenced. This workflow was repeated twice using the same PEG-concentrated sample, thereby providing opportunities for assessing the reproducibility of the system. The tests yielded a phage hit ratio of 0.084 – 0.11%, with corresponding statistics summarized in **Fig. 2B**. The droplet sorting process during the PRISM assay is shown in **Movie S1**, and representative examples of the confirmation of PRISM dispensed plaques/zones-of-clearing are shown in **Fig. S2**.

**Fig 2.**
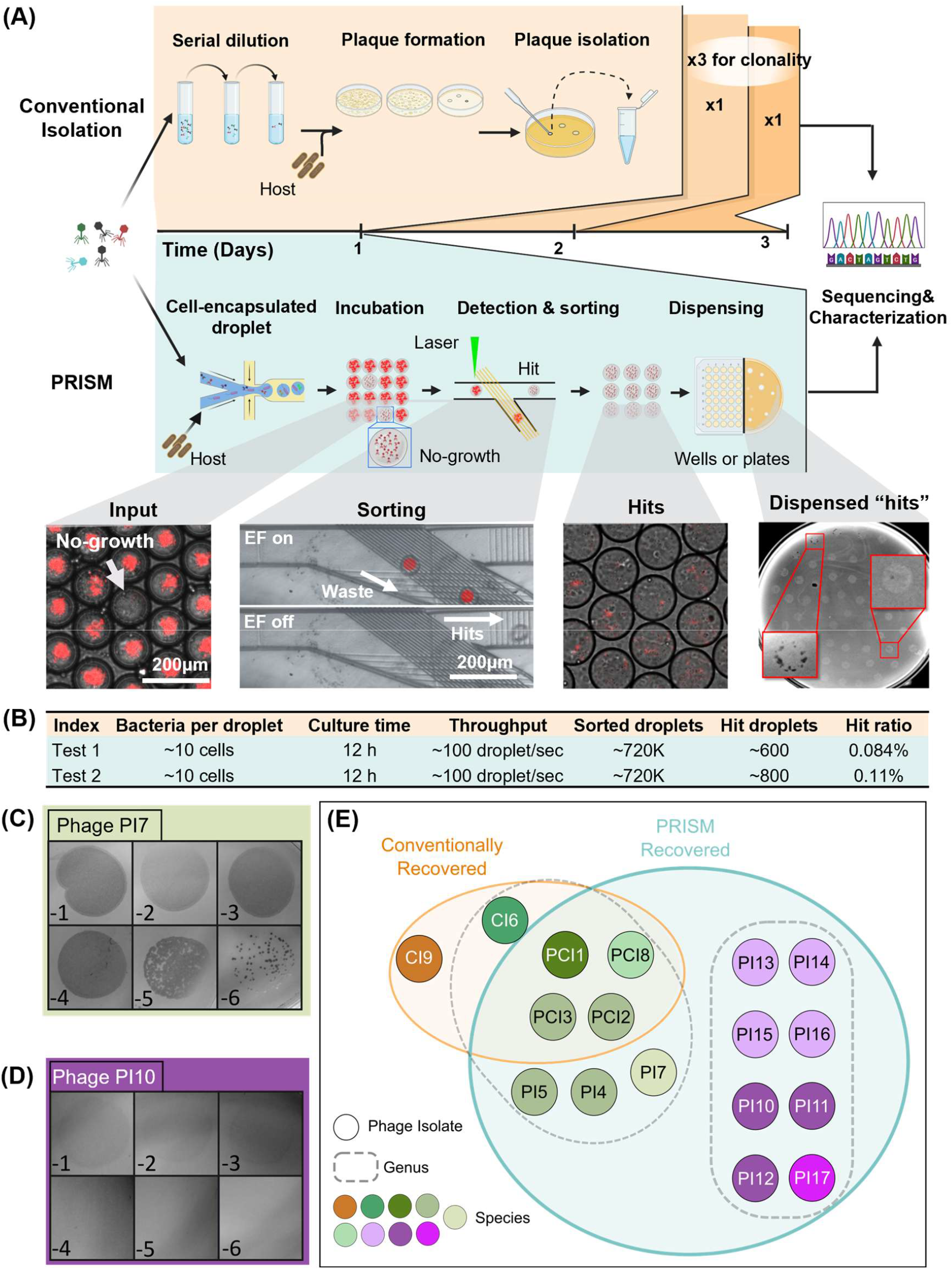
PRISM enables high-throughput screening and recovery of both plaquing and non-plaquing *Salmonella* phages. **(A)** Schematic comparison of conventional agar plate-based phage isolation versus PRISM-based droplet microfluidics workflow. Red boxes highlight plaques or clear zones. **(B)** Summary of PRISM-based screening assay results. **(C, D)** Spot assays showing 10-fold serial dilutions of recovered phages, demonstrating both **(C)** plaquing and **(D)** non-plaquing phage phenotypes. **(E)** Venn Diagram representing recovered phages taxonomically clustered based on genome alignment of sequenced phages using Progressive Mauve.

The sorted “hit” droplets from Test #2 were then closely examined. A total of 80 regions in the *Salmonella* top-agar lawns showed possible evidence of phage activity (**Fig. 2A Dispensing)**. Of these, 24 regions were capable of further phage propagation. PCR analysis identified 11 of the isolates as matches to conventionally recovered phages. The remaining 13 were uniquely recovered by PRISM. Of those 13, 11 were successfully sequenced for further analysis. Notably, PRISM-recovered phages fell into two distinct phenotypes of plaque morphology. One morphology, represented by Phage PI7, was characterized by standard plaquing that produced distinct circular zones of clearing, which resolved into individual plaques upon serial dilution (**Fig. 2C**). The other phenotype, represented by Phage PI10, was characterized by a “non-plaquing” morphology. In this case, high dilutions of phages resulted in only faint turbidity, and further dilution did not yield individual plaques (**Fig. 2D**).

In total, the conventional and PRISM approaches recovered 6 and 15 unique phages, respectively. Importantly, 11 of the PRISM-recovered phages were unique to the PRISM approach. Notably, a group of these phages were found to display “non-plaquing” phenotype, which did not form visible plaques under standard top-agar overlay plating parameters, and was presumably the reason that they could not be identified by conventional strategies (**Fig. 2E**).

### Taxonomic diversity recovered by PRISM vs. conventional isolation

Sequencing of the unique phages and genomic comparison indicated that, cumulatively, the PRISM and conventional isolation approaches recovered 9 distinct taxonomic species, using 95% nucleotide identity as a cutoff^20^. The conventional approach recovered phages representing five distinct species, while PRISM recovered representatives from seven distinct species (**Figs. 2E, S3A**). Notably, the PRISM-recovered phages include three groups of related “non-plaquing” species, which were not observed using the conventional approach.

To assign a qualitative metric of relative novelty to the isolated phages and determine if non-plaquing phage species were globally recovered less frequently compared to their plaquing counterparts, a representative genome from each species was compared to deposited phages in the nr/nt NCBI database (**Fig. S3A**). Given that *Salmonella* phages are highly studied and well represented in the NCBI database, we expected that the number of returned hits would be an indicator of the relative frequency with which a particular phage type is recovered in the field of phage research, i.e. its novelty. To return even distantly related phages from the database, a 70% nucleotide identity and query coverage was selected as a cutoff, in general, it was found that species which were recovered by both the conventional approach and the PRISM approach were more highly represented in the database, returning approximately 110 similar phage hits. Conversely, PRISM returned approximately half as many hits, indicating they are less frequently recovered.

To explore the genomic structures and similarities of the recovered phages, a representative genome from each species was annotated and compared to view overall synteny and genomic arrangement (**Fig. S3B**). Phages CI6, PCI8, PCI1, PI7, and PC13 were selected as representatives of their respective species within the same genus for comparative analysis. These genomes are approximately 157 kb in length and share characteristic features of T4-like phages. They also exhibit 85–90% sequence identity with the well-annotated *Salmonella* phage Mooltan^21^. The species sharing the non-plaquing phenotypes, represented by phages PI13, PI10, and PI17, were highly syntenic, having genomes of ∼60 kb and ∼93% nucleotide similarity to the well-studied model *Salmonella* phage Chi^22^. Finally, the most novel phage species recovered, represented by phage CI9, was approximately 55 kb in length and ∼80% nucleotide identity to *Salmonella* phage 9NA^23^. Together, these results indicate that PRISM recovered multiple unrelated phage types from a single sample, supporting its recovery of high levels of phage diversity.

### Plaque-independent phage titering using PRISM

The original motivation behind PRISM was to broaden the diversity of detectable phages by reducing dependence on plaque formation. Interestingly, the non-plaquing PRISM phages identified in this study were discovered coincidentally. While *Salmonella* species typically support robust and easily countable plaque formation, we have observed that some phage-host systems fail to produce visible plaques reliably, despite extensive optimization efforts. This inconsistency, along with our recovery of non-plaquing PRISM phages, promoted us to explore PRISM as a plaque-independent method of phage quantification (**Fig. 3A**). In this context, the infection rate was defined as the ratio of total inhibited droplets to total screened droplets. Based on Poisson distribution, we have infection rate (droplet containing at least one phage):

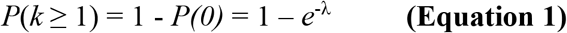

**Fig. 3.**
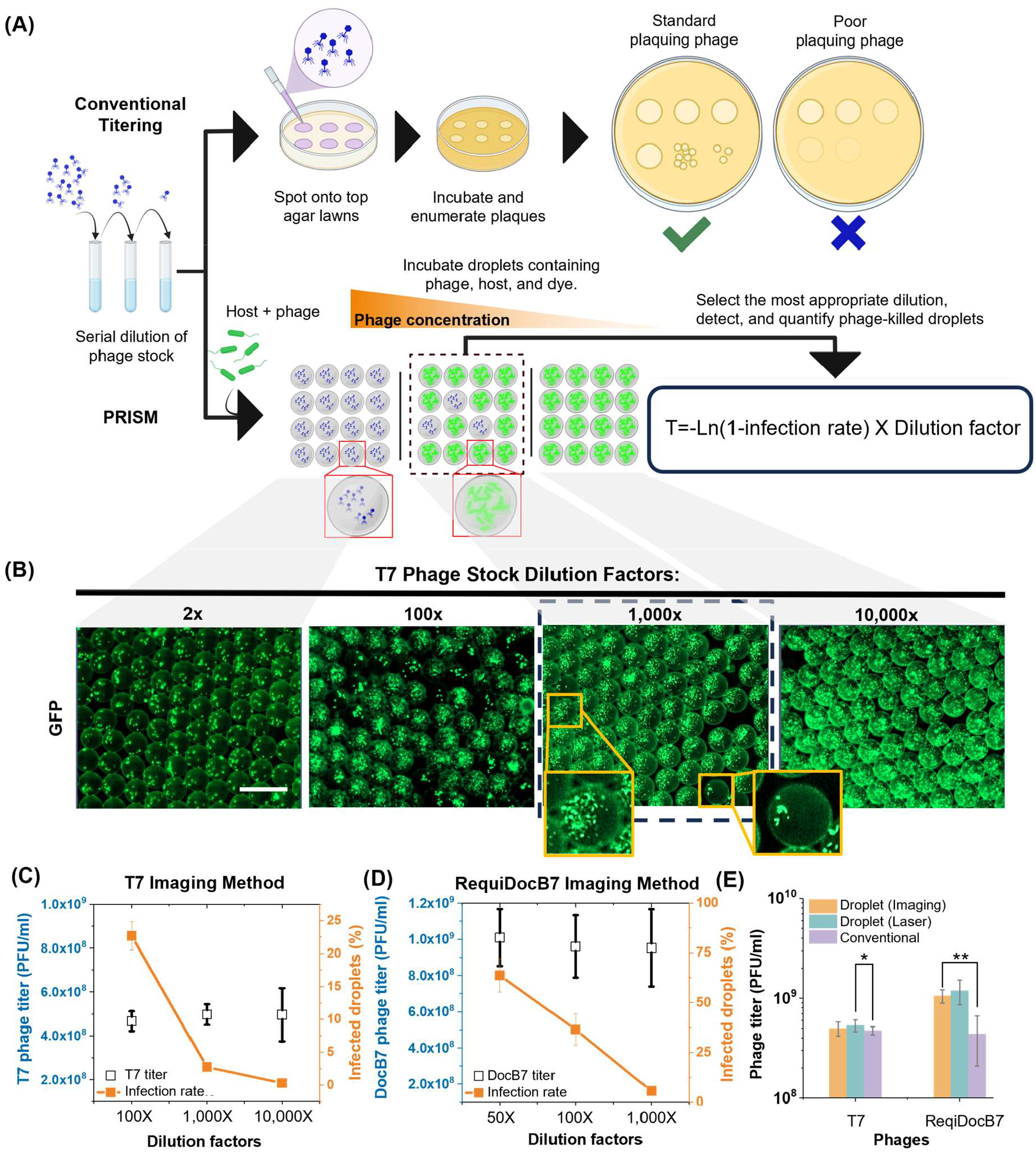
Plating-independent system for high-accuracy phage tittering. **(A)** Conventional (top row) and PRISM-based (bottom row) phage tittering. **(B)** Phage T7 co-encapsulated with *E. coli* at different dilution factors (images were taken after 12 h incubation, Scale bar: 50 µm). **(C–D)** Infection rates (orange) and calculated titers (blue) of T7 **(C)** and ReqiDocB7 **(D)** phages using fluorescent imaging-based droplet analysis. **(E)** Plaque forming (T7) and non-plaque-forming (ReqiDocB7) phage tittering using PRISM and the conventional method (one-way ANOVA test, “^*^”, *P* = 0.139, and “^**^”, *P* = 0.022, N = 3).

Where λ is the average number of phage in each droplet. Based on equation 1, we have Equation 2:

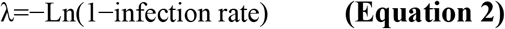

Therefore, phage titer can be calculated through the infection rate, dilution factor, and Poisson distribution. Specifically, the titer (T) is determined by the following equation 3:

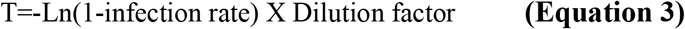

To validate the accuracy of the PRISM-based titering method, we first tested it using the well-characterized reliable plaque-forming phage T7 and its *E. coli* host. The infection rate of T7 phages co-encapsulated with *E. coli K12* at different dilution factors was used for detection and phage quantification (**Fig. 3B & Fig. S5**). We directly compared phage titers determined by conventional plaque assays with those obtained using PRISM via both fluorescent microscopy droplet imaging and high-throughput laser-based detection methods (**Fig. 3C and 3E**). The droplet imaging method involved taking multiple micrographs and subsequent image processing to identify phage inhibited hit droplets. Two methods of PRISM-based titering were evaluated to compare their relative accuracy and necessity of high-throughput detection. Regardless of phage dilution factor and subsequent infection rate, the total percentage of killed droplets (Fig. 3C, orange-line), similar titer results were achieved used for calculation. Conventional plaque-based and PRISM-based titering of T7 were largely in agreement. Notably, laser-based detection enabled more convenient (faster readout) and more scalable analysis. The higher phage titer determined by PRISM was consistent with trends of plaque-independent liquid-based methods^24-26^, and confirmed that PRISM can yield reliable and accurate titer measurements.

Having validated our PRISM titering approach, we used an environmental wild-type bacterium *Rhodococcus equi* and the previously isolated phage *Rhodococcus* phage ReqiDocB7^27^ to extend this use case. Due to the poor-plaque formation (faint plaques that are difficult to visualize) associated with phage ReqiDocB7 on *R. equi* lawn (**Fig. S4**), plaque based titering of ReqiDocB7 lysates by traditional top-agar overlay cannot be reliably used. Therefore, we sought to establish PRISM’s ability to accurately quantify the titer of RequiDocB7. Conventional titering estimated RequiDocB7 to be 3.76×10^8^ PFU/ml. However, this method exhibited high variability due to poor plaque formation (as can be seen by the large error bars in **Fig. 3E)**. In contrast, PRISM-based titering using either microscopy or laser detection yielded a more consistent result, estimating the titer at approximately 1×10^9^ PFU/mL, about twice as high, with significantly reduced variation (**Fig. 3E**). This finding demonstrates that the PRISM-based detection of phage-inhibited droplets could achieve a more accurate determination of poor plaquing phage titer when conventional titer determination is unreliable. Together, these results demonstrate the PRISM-based approach enables accurate quantification of phages even for phage-host pairs that do not support plaque formation, thereby extending its utility beyond phage isolation.

### PRISM determination of bacterial phage resistance and phage lifestyle

In addition to phage isolation and quantification, we used PRISM to characterize the phage resistance frequency of bacteria and phage lifestyle (**Fig. 4A bottom**). In the conventional approach, a plate is seeded with a high concentration of phage and host culture is spread on top, followed by incubation to identify and enumerate surviving colonies. Surviving colonies become insensitive to phage via bacterial mutations that confer phage resistance (e.g., by loss of the phage receptor), or by formation of lysogens, in which the phage integrates into the host genome and renders it immune to further phage infection. Adapting this concept to PRISM, phages were co-encapsulated with the host at a high multiplicity of infection (MOI), incubated, and screened for rare droplets containing high-density bacterial growth, indicating resistance to the applied phage or the formation of lysogens, using laser-based droplet fluorescence detection. This approach enabled resistant variants to be observed in droplets (**Fig. 4B**). For validation of the phage resistance or lysogenization frequency determination, we used the well characterized *E. coli* phage T7 (lytic phage lifestyle) and lambda (temperate phage lifestyle). By quantifying the number of droplets showing robust host cell growth, the total number of droplets screened, and starting number of bacterial cells per droplet, the frequency of phage resistance in the population was determined. For comparison, an aliquot of the same culture was simultaneously assayed for resistance frequency by conventional means^12,13^ (**Fig. 4A top**). The frequency of phage resistance for phage T7 determined by PRISM was one in 4.4×10^5^ bacterial cells and, in support of this value, the conventional approach estimated resistance frequency to be one in 4.2×10^5^ bacterial cells (**Fig. 4C**). These values were also consistent with the reported frequency of phage T7 resistance^28^.

**Fig. 4.**
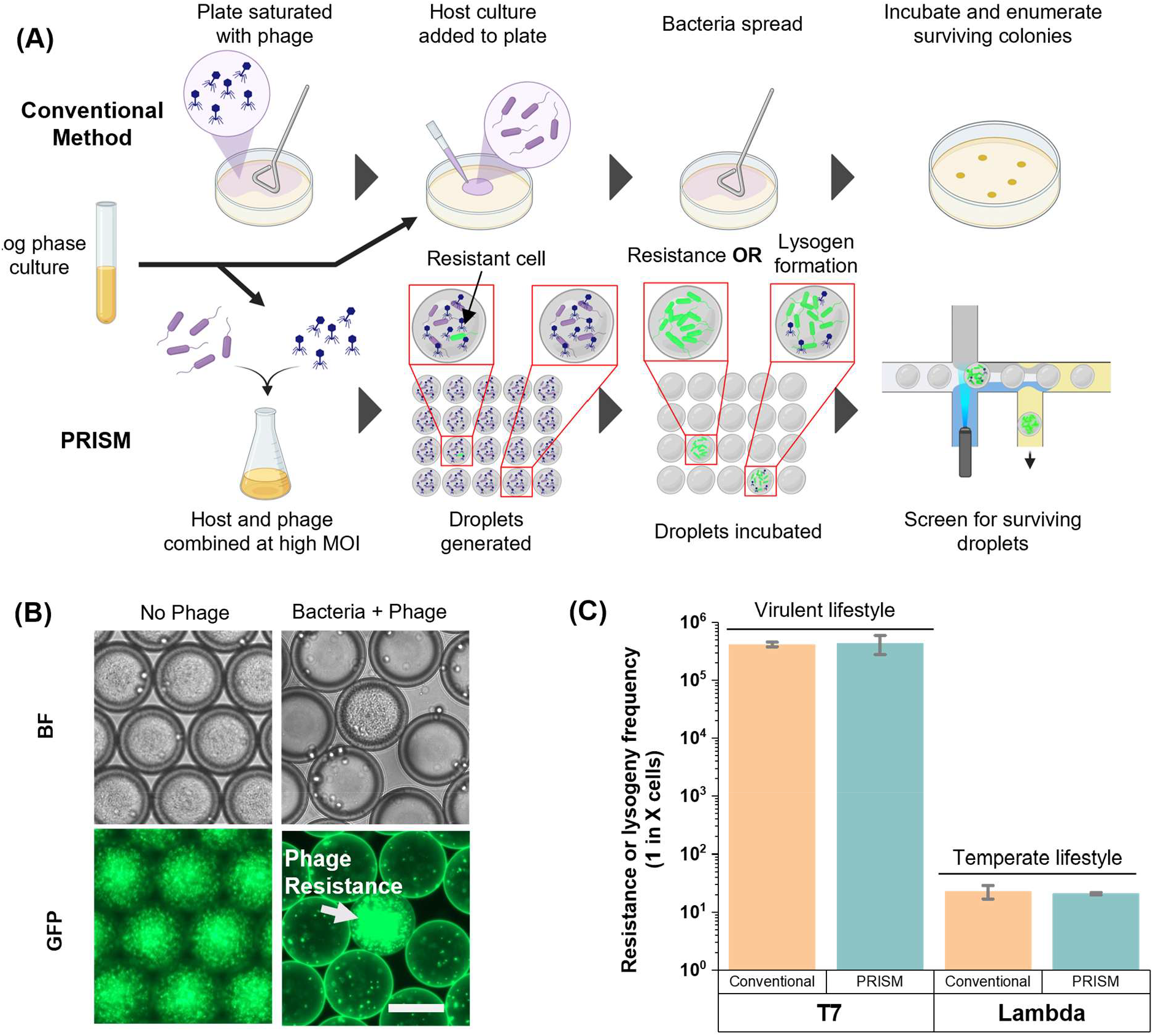
PRISM-based plating-independent system that can determine phage resistance frequency, recover phage-resistant bacterial mutants, and predict phage lifestyle. **(A)** Quantification of phage-resistant mutants using conventional and PRISM methods. **(B)** Example of rare droplet containing T7 phage-resistant *E. coli* when infected at high MOI (MOI = 100), scale bar: 50 µm. **(C)** Frequency of T7 resistance as determined by conventional or PRISM approaches, and determination of lysogen formation in droplets utilizing phage lambda. Discrepancy in frequency of surviving droplets indicates lifestyle probability. N=3

In addition to phage resistance frequency, we tested the hypothesis that PRISM enables rapid prediction of temperate and virulent phage lifestyles, which are key features for therapeutic phage selection^1,7^. Utilizing the model temperate phage lambda, lysogenized cells were recovered at rates corresponding to one in 23 cells when using the conventional method and one in 21 cells when using the PRISM approach (**Fig. 4C**).

While the lysis–lysogeny decision remains incompletely understood for many temperate phages, studies have shown that higher MOI often correlates with increased rates of lysogeny for several well-characterized phage repressor systems^11^. In other cases, lysogeny occurs at a relatively constant rate that is significantly higher than the frequency of mutational resistance^10^. Therefore, it is expected that temperate phages assayed at high MOI in the PRISM system will always produce surviving lysogenized droplets at a frequency considerably greater than that of the mutational resistance frequency to a phage, allowing identification of temperate phage (**Fig. 4C**). Together, these findings demonstrate that in addition to phage isolation, the PRISM system has additional utility for phage characterization in a plating-independent manner.

## Discussion

While recent studies have begun investigating phage isolation using microfluidic platforms^29 30^, PRISM represents the first system for efficient phage isolation and characterization. Furthermore, benchmarking of PRISM to conventional approaches from the same input sample established working parameters for the developed technology. PRISM has the potential to streamline phage isolation and characterization workflows by removing the need for time-consuming subculturing to ensure clonality, shortening the overall process by multiple days. In fact, phages could be isolated in under 24 hours, at least 3× faster than traditional methods, and with significantly reduced labor and materials cost.

The PRISM system demonstrates inherently higher phage–host contact sensitivity compared to conventional agar plate methods^31-33^. In a confined picoliter-scale droplet, a single phage following Brownian motion dynamics remains within micrometers of its bacterial targets, drastically increasing the likelihood of successful encounter, adsorption, and infection of its host^31,32^, as well as the subsequent detection of these events via fluorescence-based cell enumeration. In contrast, in conventional plaque-based assays, phages must rely on passive diffusion through a semi-solid medium, where spatial separation, limited mobility, and inactivation on cell debris, amongst other factors, reduce the probability of successful host cell infection. As the eventual formation of a visible plaque requires multiple successive rounds of infection, these limitations of semi-solid media assays are compounded, particularly for phages that do not propagate efficiently or exhibit poor diffusion capabilities. These fundamental differences in liquid vs solid-state encounter dynamics contribute to our observations regarding more accurate estimated titer and phage infectivity. First, droplet-based phage titers tend to be higher than those determined by plaque assays, especially for poor-plaquing or non-plaquing phages, as demonstrated by *Rhodococcus* phage ReqiDocB7 **(Fig. 3**). This observation is consistent with trends reported by other plaque-independent enumeration approaches, such as most probable number (MPN), which generally report higher titers than plaque assays where plaque forming units can be considered closer to the minimum number of viable particles in a solution^24-26^. The greater discrepancy between titer values obtained by PRISM and the plaque assay for ReqiDocB7 compared to T7 can likely be attributed to the high adsorption rate and “hyper virulent” nature of T7^34^, whose infection dynamics make it better suited for fairly accurate enumeration via plaque-assay. These features of T7, as well as those of phage Meme, likely also explain their single-particle infection rates closely approximating the predicted theoretical values (**Fig. 1D** and **1E)**. The forced spatial proximity and repeated contact opportunities in droplets are more likely to overcome physical and stochastic failure modes common in bulk culture or solid media, leading to improved detection of phage activity.

We successfully isolated multiple novel phages from the environment via PRISM. Many of these PRISM-recovered phages were shown to be “non-plaquing” under standard phage plating conditions used in this study (**Fig. 3**), indicating they otherwise would have gone undetected in the absence of our PRISM system. We hypothesize that the amplification of the phage within the microdroplet and the subsequent deposition of hundreds or thousands of clonal phage particles onto the top-agar lawn facilitated disruption of the bacterial lawn and their subsequent isolation. Conversely, the deposition of a single phage particle from a non-plaquing phage would likely have little impact on the bacterial lawn, and as such were not recovered by conventional isolation methods. Despite this outcome, the “non-plaquing” phages described here had nucleotide similarity to numerous phages deposited in the NCBI database, albeit fewer hits than their plaquing counterparts recovered in this study. Given plaque formation is dependent on a variety of factors such as phage adsorption rate, host growth rate, and diffusion rates, amongst other factors^35^, the isolation of similar phages to the “non-plaquing” ones described here can be attributed to the use of an alternate *Salmonella* strains or altered plating conditions. Nevertheless, this further exemplifies the need for a more universal phage isolation methodology that is less dependent on artefacts affecting plaque formation. Having recovered a greater taxonomic diversity, PRISM therefore has the potential to discover otherwise untapped phage reservoirs which could be sources of novel “viral dark matter” of great biomedical value.

Finally, we further advanced phage-host interaction characterization by quantifying the frequency of bacterial resistance to phage infection and implementing a rapid and robust approach for predicting phage lifestyle. These two parameters—resistance frequency and lifestyle classification—represent critical determinants in evaluating the therapeutic potential and applicability of newly recovered phages^7,12,13,36,37^. Accurate determination of resistance prevalence enables the assessment of selective pressures imposed by individual phages and lends insight into the durability of their efficacy in clinical or environmental contexts. Simultaneously, rapid lifestyle prediction—differentiating lytic from lysogenic phages—is essential, as only obligately lytic phages are generally desirable for antimicrobial applications due to their ability to efficiently clear bacterial populations without integrating into the host genome and rendering the host insensitive to the phage. Importantly, our ability to isolate and analyze libraries of phage-resistant bacterial mutants offers a powerful experimental framework for elucidating bacterial defense mechanisms and identifying molecular targets of resistance. These outcomes, in turn, will facilitate the rational design and optimization of therapeutic phage cocktails with improved efficacy and minimized resistance emergence.

## Conclusion

PRISM is a plating-independent, droplet microfluidics-based platform that enables rapid and high-throughput isolation, quantification, and characterization of bacteriophages. By compartmentalizing phage-host interactions in picoliter-volume water-in-oil droplets and using fluorescence-based droplet detection and sorting, PRISM accelerates the workflow, reduces material costs, and captures a broader spectrum of phage diversity. In head-to-head comparison using the same input samples, PRISM recovered more unique phages than conventional methods, including several that failed to form plaques on conventional soft-agar overlays but exhibited clear lytic activity. This ability to access rare or novel viral types supports the platform’s potential to recover novel viral “dark matter”. The platform further enables accurate measurement of phage titer and resistance frequency in a streamlined, agar-free format. Notably, PRISM outperforms conventional assays in titering poor-plaquing phages, offering higher precision where plaque-based methods may yield only rough estimates. Altogether, PRISM modernizes phage biology by combining speed, resolution, and universality in a single microfluidic system, with ongoing development aimed at expanding its application to diverse hosts and complex microbial samples.

## Methods

### Bacterial strains and phage propagation

*Escherichia coli* (MG1655), *Salmonella enterica* serovar *typhimurium* (ATCC 53648), and a clinical *Rhodococcus equi* (+103) strains were cultured in their preferred medium, LB (Luria-Bertani), TSB (Tryptic Soy Broth), or BHI (Brain Heart Infusion), or plated on their respective medias supplemented with agar. A chromosomal copy of RFP was previously introduced to *S*. Typhimurium. Previously isolated phages or model bacteriophages were propagated on their respective hosts by the top agar overlay technique^8^ containing 0.5% agar or 0.25% agar when necessary and supplemented with 5 mM Ca/Mg.

### PRISM chip fabrication

The PRISM droplet microfluidic system fabrication steps are similar to the method published previously^14^. In brief, a two-photon polymerization instrument was used to fabricate the master mold having sloped structures for the microfluidic device (2PP; Nanoscribe Photonics Professional GT, IP-Q photoresist). Electrode patterns were created on the glass substrate by first depositing a Ti/Au layer (20 nm/200 nm) using electron-beam evaporation, followed by photolithography and metal etching. The microfluidic channels were replicas from the master mold in polydimethyl siloxane (PDMS) using conventional soft lithography techniques.

### Generation of phage-bacteria encapsulated droplets

Bacterial cultures were prepared by diluting an overnight saturated culture 1:500 v/v in tryptic soy broth (TSB; Becton, Dickinson) and incubating at 37°C and 250 rpm for 2 h. The resulting culture was then directly diluted to an OD_600_ of 0.1 (measured using a Nanodrop spectrophotometer, Thermo Scientific). Fluorescent cell stain SynaptoGreen™ C4 (Biotium) was added to the bacterial suspension to enable fluorescence enumeration of cells in droplets. When required, phages targeting the bacterial strain were added to the same aqueous phase at varying multiplicities of infection (MOIs), depending on the experimental design.

Phase-bacteria encapsulated water-in-oil emulsion droplets were generated using the PRISM device having a flow focusing droplet generator geometry (channel cross-section: 50 μm × 50 μm). The aqueous phase (bacteria and phage in TSB) was introduced at a flow rate of 300 μL h-1 using a syringe pump (KDS Legto 200™), while the continuous oil phase Novec 7500 (3M) containing 2% (w/w) PFPE-PEG surfactant (PicoSurf 1, Sphere Fluidics) was introduced at 500 μL h-1 using a syringe pump (Legato Systems). All fluids were delivered via Tygon tubing (Saint-Gobain, ID 0.25 mm) and connected to the PRISM chip immediately after surface treatment with 1% trichloro(1H,1H,2H,2H-perfluorooctyl) silane (Merck) in Novec 7500 for 1 min to minimize droplet adhesion. Droplets (∼50 μm in diameter, ∼65 pL in volume) encapsulated approximately 10–20 bacterial cells each and were produced at a rate of ∼100 droplets per second.

### Incubation, droplet counting, and sorting

After generation, droplets were incubated at 37°C in a microfluidic chamber for up to 24 h to allow for phage infection, bacterial lysis, or continued bacterial growth. In this system, successful phage infection leads to bacterial lysis, while uninfected or phage-resistant bacteria proliferates, producing a strong fluorescent signal within the droplet. A flow-through droplet fluorescence detection setup (**Fig. S6A**) was used to identify and count high-fluorescence “hit” droplets for laser-based detection. Fluorescence signals were processed using an FPGA I/O data acquisition card (PCIe-7842, National Instruments) and a custom LabVIEW™ program (**Fig. S6B**). These signals enabled quantification of both total and high-signal droplets for downstream statistical analysis. The detection system was integrated with a feedback-controlled droplet sorting mechanism that activated electrodes to apply dielectrophoretic force to droplets. Correct-size droplets exceeding a defined fluorescence threshold were selectively sorted^14^.

### Operation of the PRISM droplet sorting function

All droplet populations were reintroduced into the PRISM droplet sorting device at a flow rate of 35 μL/h, resulting in a throughput of approximately 30 Hz. The spacing fluid was flowed at 500 μL/h, with Bias 1 and Bias 2 set to 800 μL/h and 1000 μL/h, respectively. Flow rates were controlled using syringe pumps (KD Scientific Legato series). A 10 kHz, 60 peak-to-peak voltage (Vpp) Vpp square wave was applied to the first set of electrodes via a waveform generator (Rigol DG4102). As bright fluorescence droplets pass through the detection region, a feedback loop triggered the second set of electrodes with a 10 kHz, 40 Vpp square wave for 25 ms to perform droplet sorting. The Vpp and sorting duration were adjusted depending on the droplet throughput requirements.

### Conventional and PRISM-based phage isolation

Wastewater samples were collected from three different wastewater treatment plants in the surrounding College Station, Texas area. Samples were centrifuged at 8,000 x g for 10 min before being sterilized by passage through a 0.22 μm filter and stored at 4ºC until use. To generate a concentrated unenriched phage sample, 60 ml of each sample were pooled and precipitated by addition of PEG-8000 and NaCl to a final concentration of 10% and 1M, respectively, before overnight incubation at 4ºC. This sample was then spun at 10,000 x g for 10 min, and the resulting pellet resuspended in a minimum volume of approximately 1 ml of lambda phage diluent (25 mM Tris-HCl, 100 mM NaCl, 8 mM MgSO4, 0.01% gelatin). For conventional enrichment-based phage isolation, 10 ml of individual sterile wastewater supernatant was added to 40 ml of TSB and inoculated with 100 μl of an overnight (16 – 18 hr) host culture before being incubated overnight at 37ºC with aeration. After overnight incubation, 10 ml of this enrichment culture was centrifuged at 10,000 x g for 10 min at 4ºC before being filter-sterilized through a 0.22 μm syringe filter. This enrichment supernatant was then screened on top-agar overlays for plaque formation or zones of clearing. For conventional isolation, morphologically distinct plaques within either enriched samples or PEG-concentrated samples were subcultured for purity and high titer stock lysates generated. For PRISM-based phage isolation, the PEG-concentrated environmental sample was added to TSB containing *Salmonella* host and encapsulated to a target of ∼20 cells per droplet. After incubation, phage “hit” droplets were detected based on lack of RFP signal, sorted, and combined with empty droplets to ensure sufficient distribution of “hits” such that the clonality of phage population within an individual droplet is maintained when deposited onto top-agar overlays seeded with *Salmonella*. Plaques or burn through present in deposited droplet zones were collected for further analysis and/or used to for the generation of high titer lysates.

### Droplet Dispensing and Hit Validation

Hit droplets were dispensed using previously published single-droplet-resolution dispensing method^38^, which employs blank droplets as spacers to isolate and deliver individual “hit” droplets with high precision. This strategy ensures physical separation between droplets, preventing cross-contamination without the need for complex feedback control. The “hit” droplets were dispensed onto top-agar overlays seeded with host bacteria of interest. Dispensed “hit” droplets were allowed to dry before being incubated and inspected for phage activity. Droplets that caused some level of inhibition of the bacterial lawn were collected for downstream sequencing to determine the genotype of the corresponding phage.

### Conventional vs PRISM-isolated phage comparison

To discriminate novel phages isolated from the PRISM approach that were different than those identified by the conventional approach, conventionally isolated phages were first sequenced and discriminatory PCR primers designed for detection of the conventionally isolated phages (**Supplemental Table 1**). Pickates recovered from the deposited microdroplets were screened for identity to the conventional phage by plaque PCR utilizing 1 μl of pickate or 1 μl of a 1/10th dilution of pickate into H_2_O as template. Plaque PCR was conducted using 2x Q5 Mastermix with the following thermocycler conditions: 98ºC for 10 min; 30 cycles of 98ºC for 30 sec, 60ºC for 20 sec, 72ºC for 20 sec with a final extension of 72ºC for 5 min. PRISM originating pickates generating an amplicon with any of the discriminatory primer sets were determined to likely be a match for one of the conventionally isolated phages, and thus removed from further downstream propagation and analysis. Those pickates that generated no amplicon products for any of the discriminatory primer sets were presumed to be novel, and thus a high titer lysate generated, DNA extracted, and sequenced as described above.

### Phage DNA and genome analysis

DNA was extracted from high titer phage lysates as described previously using either Phenol:Chloroform-based extraction or silica DNA spin column-based isolation techniques^39^. Resulting DNA was prepared for Illumina sequencing using tag mentation based and PCR based Illumina DNA prep kit, and ultimately sequenced on an Illumina NovaSeq6000 platform by SeqCenter (Pittsburgh, PA). Resulting reads were assembled and analyzed on the Texas A&M University Center for Phage Technology instance of Galaxy^40^. After genome closure, genomes were aligned and compared at the nucleotide level by ProgressiveMauve^41^ enabling taxonomic clustering. Phage genomes from a representative of each taxonomic species identified was automatically annotated with Prokka utilizing settings^42^ with parameters selected for viruses on the European instance of Galaxy (Usegalaxy.eu). Annotated genome maps were generated via EasyFig using the setting tblastX for homology detection.

### Determination of frequency of resince by conventional and PRISM approaches

Bacterial cultures were grown to early log phase (approximately OD_600_ of 0.2 - 0.4) and the cultures were split to be assayed in parallel by the conventional and PRISM approaches to account for founder effects of phage resistant bacteria. For conventional determination, plate was flooded with 100 µl of high titer phage lysate at a concentration >10^9^ PFU/ml for a high MOI and allowed to dry in air. Host culture (100 µl) was then applied, incubated overnight at 37ºC, followed by enumerating the surviving colonies. Additionally, host culture was serial diluted and plated to determine the total CFU/ml of the starting culture. For PRISM, phages were added to host culture at an MOI >10 before droplet encapsulation, incubated, and “hit” droplets (high fluorescent intensity) detected and sorted. Additionally, during the validation of the PRISM approach, sorted “hit” droplets were dispensed onto agar plates coated with bacteriophage and incubated. At least 10 resulting colonies were subcultured for clonality analysis, and resistance to T7 phage was confirmed by cross-streaking assays. Furthermore, sorted fluorescence-dark droplets were dispensed onto T7-coated plates, which yielded no colonies. To confirm these fluorescence-dark droplets were killed, they were also dispensed onto plates with no phages, which yielded rare colonies; however, these colonies were found to not be resistant to phage T7 and instead represented rare persister cells, which could only grow once phage T7 was removed.

## Credit Authorship Contribution Statement

**Han Zhang:** Conceptualization, Methodology, Validation, Investigation, Writing – original draft, Writing – review & editing, Data analysis and curation. **Justin Boeckman:** Conceptualization, Methodology, Validation, Investigation, Writing – original draft, Writing – review & editing, Data analysis and curation. **Rohit Gupte:** Data curation. **Ashlee Prejean:** Data curation **Jonathan Miller:** Data curation. **Alexandra Rodier:** Data curation. **Paul de Figueiredo:** Resources, Writing – review & editing, Supervision, Funding acquisition. **Mei Liu:** Resources, Writing – review & editing, Supervision, Funding acquisition. **Jason Gill:** Resources, Writing – review & editing, Supervision, Funding acquisition. **Arum Han:** Conceptualization, Resources, Writing – review & editing, Supervision, Funding acquisition.

## Acknowledgment

The project depicted was sponsored by the Defense Advanced Research Projects Agency (DARPA) cooperative agreement HR00112320006. The content of the information does not necessarily reflect the position or the policy of the government, and no official endorsement should be inferred.

