## Supplemental Material for "PRISM: A platform for illuminating viral dark matter"

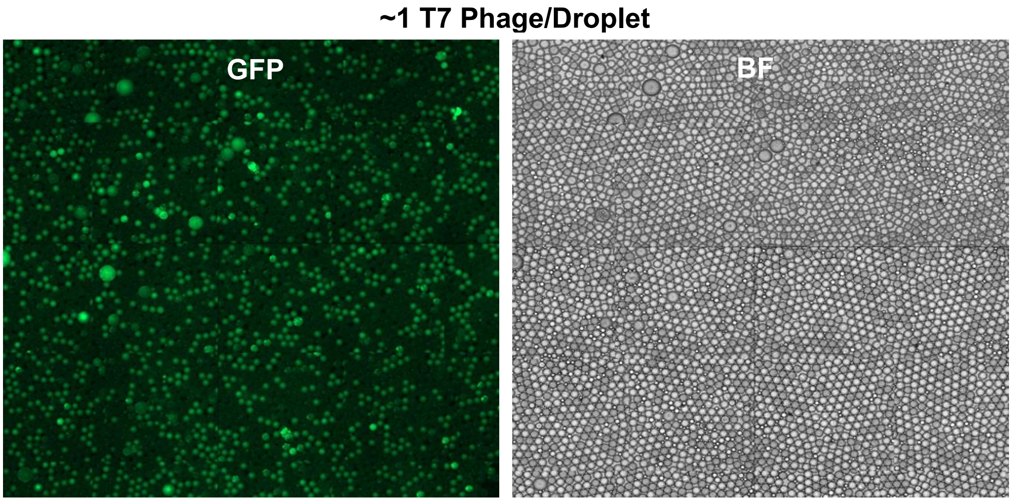


**Supplementary Figure 1. The images showing T7 phage infection of *E.coli* at 1 phage particle/ droplet.**

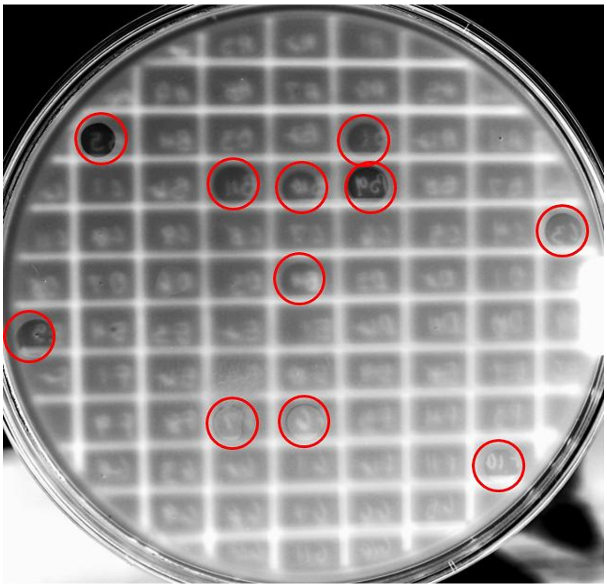

**Supplementary figure 2. Representative confirmation of PRISM plaques** Following direct dispensing of hit droplets onto top agar overlays, dispensed zones that displayed evidence of phage activity were picked from the plate and re-spotted for confirmation.

(A)

| | PCI1 | PCI2 | PCI3 | PI4 | PI5 | CI6 | PI7 | PCI8 | CI9 | PI10 | PI11 | PI12 | PI13 | PI14 | PI15 | PI16 | PI17 | Relative Novelty #nr/nt hits: |
|---|---|---|---|---|---|---|---|---|---|---|---|---|---|---|---|---|---|---|
| PCI1 | 100 | 94.39 | 92.2 | 92.08 | 91.88 | 87.52 | 79.59 | 78.9 | 1.25 | 0.07 | 0.07 | 0.07 | 0.07 | 0.1 | 0.07 | 0.07 | 0.07 | 110 |
| PCI2 | 94.39 | 100 | 97.7 | 96.73 | 97.32 | 86.33 | 82.19 | 81 | 1.22 | 0.07 | 0.07 | 0.07 | 0.07 | 0.1 | 0.07 | 0.07 | 0.07 | |
| PCI3 | 92.22 | 97.66 | 100 | 98.94 | 98.7 | 85.28 | 82.54 | 81.71 | 1.25 | 0.07 | 0.07 | 0.07 | 0.07 | 0.1 | 0.07 | 0.07 | 0.07 | |
| PI4 | 92.08 | 96.73 | 98.9 | 100 | 98.79 | 85.23 | 82.55 | 81.58 | 1.25 | 0.07 | 0.07 | 0.07 | 0.07 | 0.1 | 0.07 | 0.07 | 0.07 | |
| PI5 | 91.88 | 97.32 | 98.7 | 98.79 | 100 | 85.43 | 82.71 | 81.8 | 1.25 | 0.07 | 0.07 | 0.07 | 0.07 | 0.1 | 0.07 | 0.07 | 0.07 | 111 |
| CI6 | 87.52 | 86.33 | 85.3 | 85.23 | 85.43 | 100 | 81.51 | 80.59 | 1.57 | 0.07 | 0.07 | 0.07 | 0.07 | 0.1 | 0.07 | 0.07 | 0.07 | 111 |
| PI7 | 79.59 | 82.19 | 82.5 | 82.55 | 82.71 | 81.51 | 100 | 93.21 | 1.06 | 0.07 | 0.07 | 0.07 | 0.07 | 0.1 | 0.07 | 0.07 | 0.07 | 111 |
| PCI8 | 78.9 | 81 | 81.7 | 81.58 | 81.8 | 80.59 | 93.21 | 100 | 1.07 | 0.07 | 0.07 | 0.07 | 0.07 | 0.1 | 0.07 | 0.07 | 0.07 | 110 |
| CI9 | 1.25 | 1.22 | 1.25 | 1.25 | 1.25 | 1.57 | 1.06 | 1.07 | 100 | 0 | 0 | 0 | 0 | 0 | 0 | 0 | 0 | 5 |
| PI10 | 0.07 | 0.07 | 0.07 | 0.07 | 0.07 | 0.07 | 0.07 | 0.07 | 0 | 100 | 99.76 | 99.68 | 92.5 | 92.49 | 92.46 | 91.88 | 92.22 | |
| PI11 | 0.07 | 0.07 | 0.07 | 0.07 | 0.07 | 0.07 | 0.07 | 0.07 | 0 | 99.76 | 100 | 99.7 | 92.5 | 92.49 | 92.47 | 91.88 | 92.22 | |
| PI12 | 0.07 | 0.07 | 0.07 | 0.07 | 0.07 | 0.07 | 0.07 | 0.07 | 0 | 99.68 | 99.7 | 100 | 92.45 | 92.44 | 92.48 | 91.82 | 92.23 | 54 |
| PI13 | 0.07 | 0.07 | 0.07 | 0.07 | 0.07 | 0.07 | 0.07 | 0.07 | 0 | 92.5 | 92.5 | 92.45 | 100 | 99.74 | 99.73 | 98.47 | 92.51 | |
| PI14 | 0.1 | 0.1 | 0.1 | 0.1 | 0.1 | 0.1 | 0.1 | 0.1 | 0 | 92.49 | 92.49 | 92.44 | 99.74 | 100 | 99.7 | 98.46 | 92.5 | |
| PI15 | 0.07 | 0.07 | 0.07 | 0.07 | 0.07 | 0.07 | 0.07 | 0.07 | 0 | 92.46 | 92.47 | 92.48 | 99.73 | 99.7 | 100 | 98.42 | 92.59 | |
| PI16 | 0.07 | 0.07 | 0.07 | 0.07 | 0.07 | 0.07 | 0.07 | 0.07 | 0 | 91.88 | 91.88 | 91.82 | 98.47 | 98.46 | 98.42 | 100 | 92.49 | 54 |
| PI17 | 0.07 | 0.07 | 0.07 | 0.07 | 0.07 | 0.07 | 0.07 | 0.07 | 0 | 92.22 | 92.22 | 92.23 | 92.51 | 92.5 | 92.59 | 92.49 | 100 | 54 |

**Percent Identity**

<75
75
85
95
100

Prism Isolated
Conventionally Isolated
Prism+Conventionally Isolated

(B)

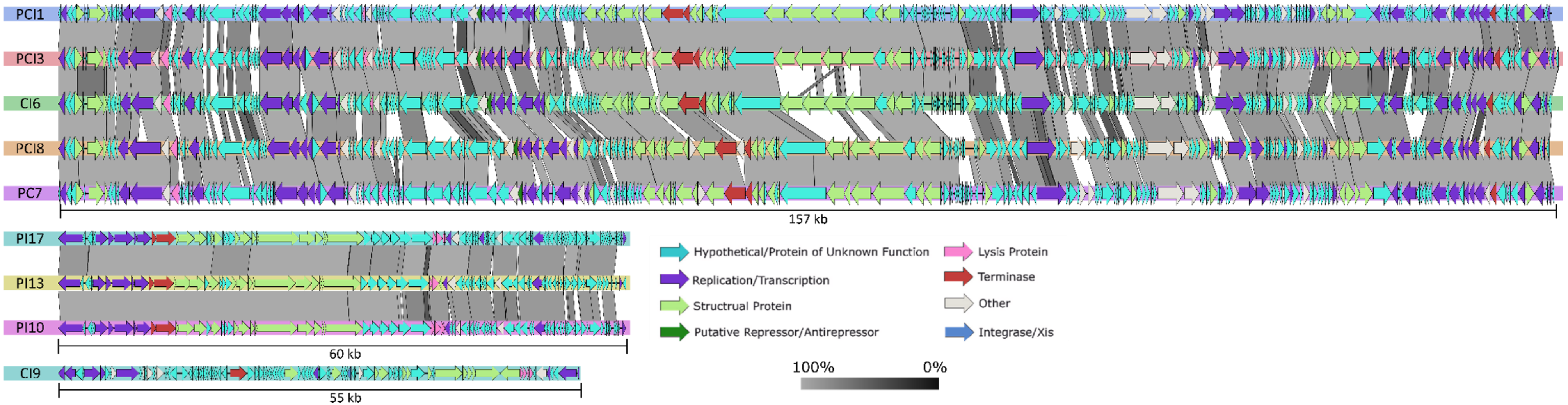


**Supplementary figure 3. Comparison of *Salmonella* phages recovered by both approaches and their relative novelty A)** Sequenced phage genome assemblies aligned and compared by Progressive Mauve and percent nucleotide identity calculated by DICE. Returned hits for a representative species isolate using a 70% nucleotide ID and query coverage (Relative Novelty). Legend heat map indicates percent identity. Colored bounding boxes indicate phage clusters within the same species (>95% ID). **(B)** Genome maps and alignment of species representatives identified in this study. Grey bars indicated regions of homology by tblastx indicated by figure legend. General function of proteins indicated by color legend. Species representative color correlates with bounding boxes in **(A)**.

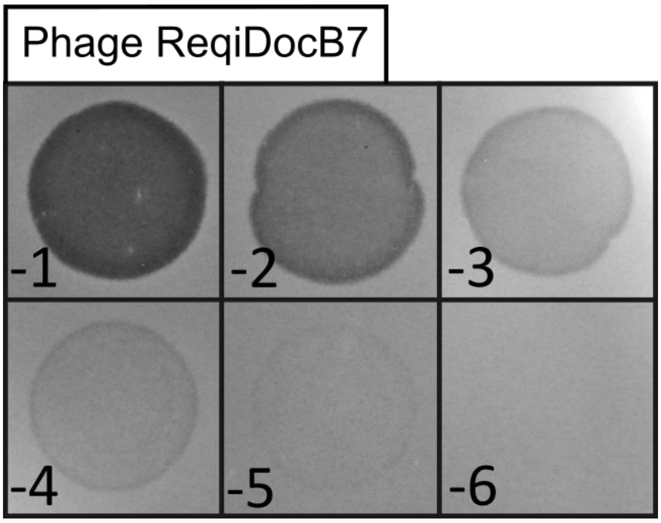


**Supplementary figure 4. The images showing non-plaque forming phenotype of phage RequiDocB7**
10-fold serial dilutions of phage RequiDocB7 spotted onto lawns of R. equi host displaying zones of burn through diluting to extremely small turbid plaques.

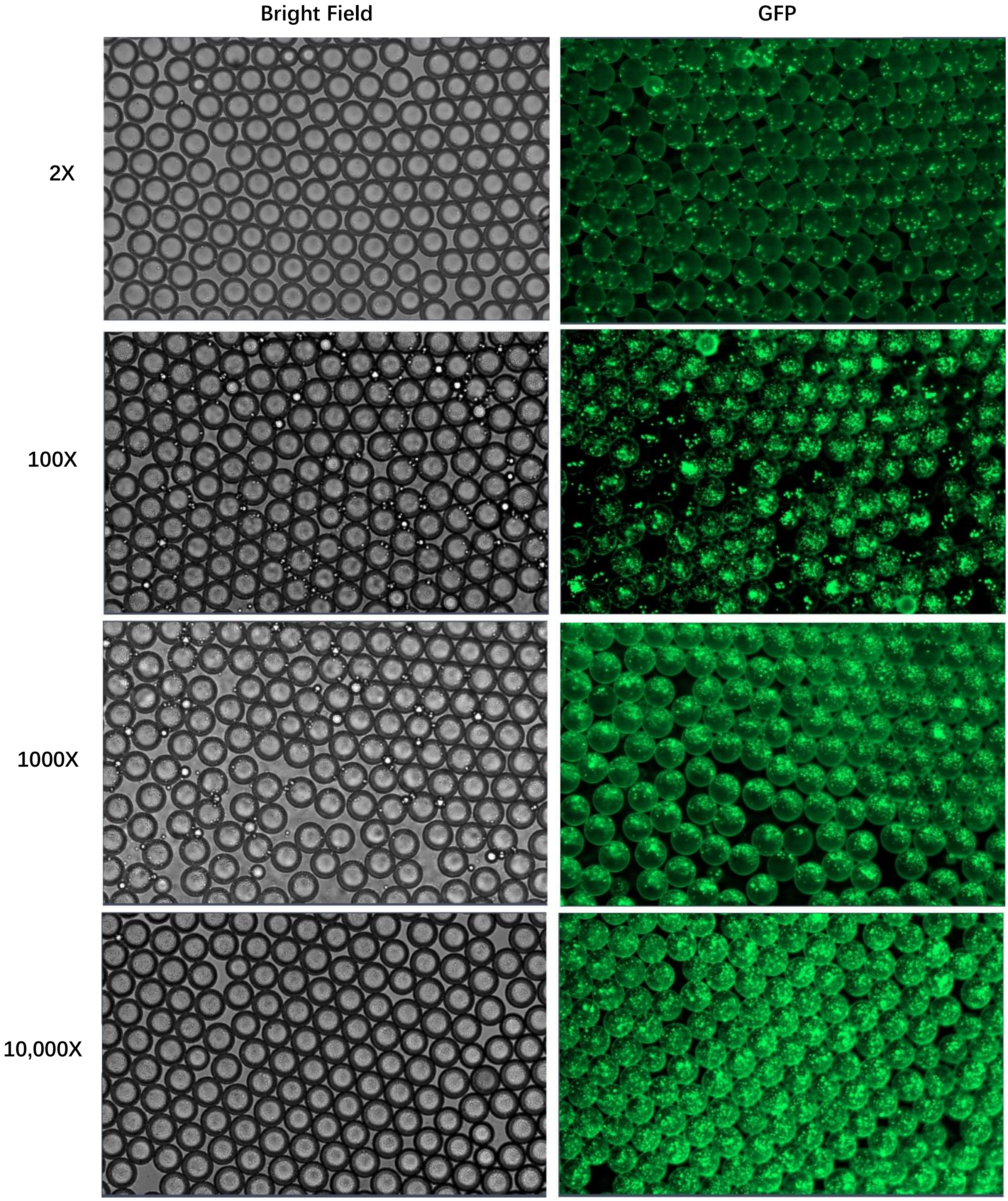


**Supplementary Figure 5. The growth inhibition rate of T7-*E.coli* at different dilution factors of T7.**

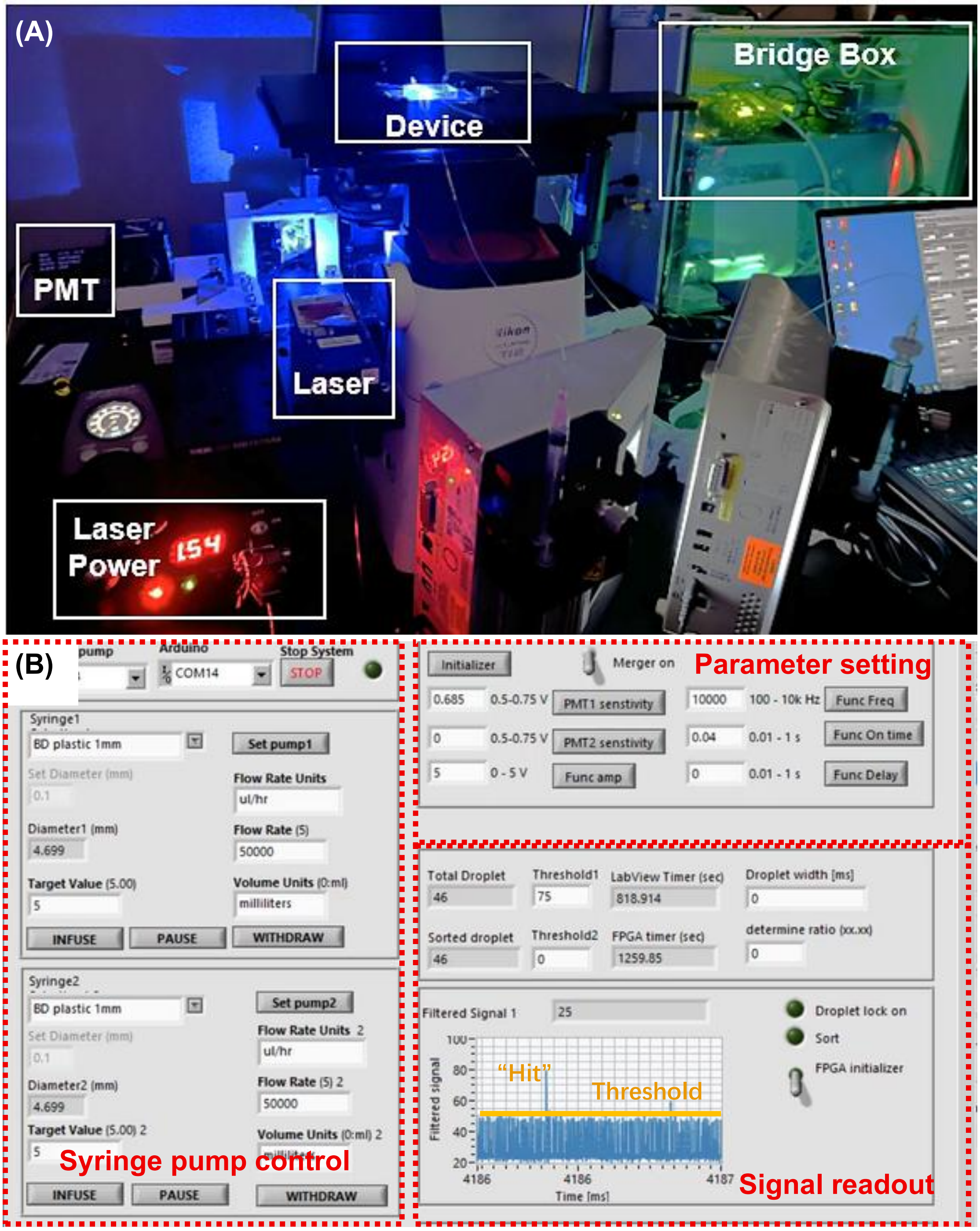


**Supplementary figure 6. The fluorescence-activated droplet sorting platform. (A)** Sorting station setup, including the electrical control sub-system (Bridge Box) and the fluorescence detection sub-system (Laser than PMT). (B) LabVIEW software user interface used for fluorescence-activated droplet sorting.

| | Source | Detection Primers F | Detection Primers R |
|---|---|---|---|
| **Phage PCI1** | TAMU Influent Enrichment | 5'-CATAATCTTCATGCCACGCC-3' | 5'-GGGATTCACTATAAAGACCGC-3' |
| **Phage PCI3** | Anahuac Influent Enrichment | 5'-GTTCGTCGGTCTGGTGATGT-3' | 5'-TTCGGAAGCCAGCTGAATTT-3' |
| **Phage CI6** | Port Arthur Activated Sludge Enrichme | 5'-GTTGCTGCCACTTTGCCTG-3' | 5'-CCCAAGCGAACTCCAGAGAA-3' |
| **Phage CI9** | PEG Concentrated Pool | 5'-CAATTGTCCCGGCTACCACT-3' | 5'-TGAACTGGGTTCACAGGACC-3' |
| **Phage PCI8** | PEG Concentrated Pool | 5'-AATGTCTTTCAGGCGAGGCA-3' | 5'-GAAGGCACCATGTGTCCGTA-3' |
| **Phage PCI2** | PEG Concentrated Pool | 5'-TTCCCCTTCCAGTAACCCGA-3' | 5'CTGGTGGTGGTATCGTCGTC-3' |
| **Additional primers for discrimination between phage PCI3 and PCI2:** | | 5'-TCAGAAGACTCGCCACCAAC-3' | 5'-CGAAACGCTGCGATGCAATA-3' |

**Supplementary table 1. Primers designed based of conventionally isolated phage sequences for deduplication and detection** original isolation source of conventionally isolated phages, and primer sequences designed for detection and discrimination.
